# Expression of a recombinant high affinity IgG Fc receptor by engineered NK cells as a docking platform for therapeutic mAbs to target cancer cells

**DOI:** 10.1101/432849

**Authors:** Kristin M. Snyder, Robert Hullsiek, Hemant K Mishra, Daniel C. Mendez, Yunfang Li, Allison Rogich, Dan S. Kaufman, Jianming Wu, Bruce Walcheck

## Abstract

Anti-tumor mAbs are the most widely used and characterized cancer immunotherapy agents. Despite having a significant impact on some malignancies, most cancer patients respond poorly or develop resistance to this therapy. A known mechanism of action of these therapeutic mAbs is antibody-dependent cell-mediated cytotoxicity (ADCC), which is a primarily effector function of NK cells. CD16A on human NK cells has an exclusive role in binding to tumor-bound IgG antibodies. Though CD16A is a potent activating receptor, it is a low affinity FcγR and its cell surface levels can be rapidly downregulated by a proteolytic process involving ADAM17 upon NK cell activation, which are likely to limit the efficacy of tumor-targeting therapeutic mAbs in the tumor environment. We sought to enhance NK cell binding to anti-tumor mAbs by engineering these cells with a recombinant FcγR consisting of the extracellular region of CD64, the highest affinity IgG Fc receptor expressed by leukocytes, and the transmembrane and cytoplasmic regions of CD16A. This novel recombinant FcγR (CD64/16A) was expressed in the human NK cell line NK92 and in induced pluripotent stem cells from which primary NK cells were derived. CD64/16A also lacked the ADAM17 cleavage region in CD16A and it was not rapidly downregulated in expression following NK cell activation during ADCC. CD64/16A on NK cells facilitated conjugation to antibody-treated tumor cells, ADCC, and cytokine production, demonstrating functional activity by its two components. Unlike NK cells expressing CD16A, CD64/16A captured soluble therapeutic mAbs and the modified NK cells mediated tumor cell killing. Hence, CD64/16A could potentially be used as a docking platform on engineered NK cells for therapeutic mAbs and IgG Fc chimeric proteins, allowing for switchable targeting elements, and a novel cancer cellular therapy.

## Introduction

Natural killer (NK) cells are cytotoxic lymphocytes of the innate immune system that target stressed, infected, and neoplastic cells (1). In contrast to the diverse array of receptors involved in natural cytotoxicity, human NK cells mediate ADCC exclusively through the IgG Fc receptor CD16A (FcγRIIIA) (2–4). This is a potent activating receptor and its signal transduction involves the association of the transmembrane and cytoplasmic regions of CD16A with FcRγ and/or CD3ζ (4–9). Unlike other activating receptors expressed by NK cells, the cell surface levels of CD16A undergo a rapid downregulation upon NK cell activation during ADCC and by other stimuli (10–14). CD16A downregulation also occurs in the tumor environment of patients and contributes to NK cell dysfunction (15–19). A disintegrin and metalloproteinase-17 (ADAM17) expressed by NK cells plays a key role in cleaving CD16A in a *cis* manner at a specific location proximal to the cell membrane upon NK cell activation (13, 14, 20).

There are two allelic variants of CD16A that have either a phenylalanine or valine residue at position 176 (position 158 if amino acid enumeration does not include the signal sequence). The CD16A-176V variant has a higher affinity for IgG (21, 22), but CD16A-176F is the dominant allele in humans (23). Clinical analyses have revealed a positive correlation between the therapeutic efficacy of tumor-targeting therapeutic mAbs and CD16A binding affinity. Patients homozygous for the CD16A valine variant (CD16A-V/V) had an improved clinical outcome after treatment with anti-tumor mAbs compared to those who were either heterozygous (CD16A-V/F) or homozygous (CD16A-F/F) for the lower affinity FcγRIIIA isoform (as reviewed in ref. 4). These findings establish that increasing the binding affinity of CD16A for anti-tumor mAbs may lead to improved cancer cell killing.

CD64 (FcγR1) binds to monomeric IgG with 2–3 orders of magnitude higher affinity than CD16A (24–26). CD64 recognizes the same IgG isotypes as CD16A and is expressed by myeloid cells, including monocytes, macrophages, and activated neutrophils, but not NK cells (24, 26). We generated the novel recombinant receptor CD64/16A that consists of the extracellular region of human CD64, for high affinity antibody binding, and the transmembrane and intracellular regions of human CD16A for optimal signal transduction. CD64/16A also lacked the membrane proximal ADAM17 cleavage site found in CD16A. In this study, we stably expressed CD64/16A in NK92 cells, a cytotoxic human NK cell line that lacks endogenous FcγRs (27), and in induced pluripotent stem cells (iPSCs) that were then differentiated into primary NK cells. We show that this novel recombinant FcγR is functional and can capture soluble monomeric IgG therapeutic mAbs that provide targeting elements for tumor cell ADCC.

## Materials and Methods

### Antibodies

All mAbs to human hematopoietic and leukocyte phenotypic markers are described in **Table 1**. All isotype-matched negative control mAbs were purchased from BioLegend (San Diego, CA). APC-conjugated F(ab’)_2_ donkey anti-human or goat anti-mouse IgG (H+L) were purchased from Jackson ImmunoResearch Laboratories (West Grove, PA). The human IgG1 mAbs trastuzumab/Herceptin and rituximab/Rituxan, manufactured by Genentech (South San Francisco, CA), and cetuximab/Erbitux, manufactured by Bristol-Myers Squibb (Lawrence, NJ), were purchased through the University of Minnesota Boynton Pharmacy. Recombinant human L-selectin/IgG1 Fc chimera was purchased from R&D Systems (Minneapolis, MN).

**Table 1.** Antibodies.

| *Antigen* | *Clone* | *Fluorophore* | *Company* |
| --- | --- | --- | --- |
| CD56 | HCD56 | PE-CY7 | BioLegend, San Diego, CA |
| CD3 | HIT3a | PE | BioLegend |
| CD16 | 3G8 | APC | BioLegend |
| CD16 | 3G8 | none | Ancell, Bayport, MN |
| CD7 | CD7-6B7 | PE/CY5 | BioLegend |
| CD336/NKp44 | P44-8 | APC | BioLegend |
| CD335/NKp46 | 9E2 | APC | BioLegend |
| CD159a/NKG2A | Z199 | APC | Beckman Coulter, Brea, CA |
| CD314/NKG2D | 1D11 | PerCP/Cy5.5 | BioLegend |
| CD158a/KIR2DL1 | HP-MA4 | PE | BioLegend |
| CD158b1/KIR2DL2/L3 | DX27 | PE | BioLegend |
| CD158e1/KIR3DL1 | DX9 | PE | BioLegend |
| CD94 | DX22 | PE | BioLegend |
| CD64 | 10.1 | APC | BioLegend |
| CD64 | 10.1 | none | Ancell |
| CD34 | 561 | PE | BioLegend |
| CD45 | 2D1 | APC | BioLegend |
| CD43 | CD43-10G7 | APC | BioLegend |
| CD62L/L-selectin | LAM1-116 | APC | Ancell |

### Generation of human CD64/16A

Total RNA was isolated from human peripheral blood leukocytes using TRIzol total RNA isolation reagent (Invitrogen, Carlsbad, CA) and cDNA was synthesized with the SuperScript preamplification system (Invitrogen). The recombinant CD64/16A is comprised of human CD64 extracellular domain and CD16A transmembrane and cytoplasmic domains. PCR fragments for CD64 (885 bps) and CD16A (195 bps) were amplified from the generated cDNA. The PCR fragments were purified and mixed together with the forward primer 5’- CGG <u>GAA TTC</u> GGA GAC AAC ATG TGG TTC TTG ACA A-3’, the reverse primer 5’- CCG <u>GAA TTC</u> TCA TTT GTC TTG AGG GTC CTT TCT-3’ (underlined nucleotides are EcoR I sites), and Pfx50 DNA polymerase (Invitrogen) to generate the recombinant CD64/16A receptor. CD64/CD16A and CD16A cDNA (CD16A-176V variant) was inserted into the retroviral expression vector pBMN-IRES-EGFP and virus was generated for NK92 cell transduction, as previously described (14). Additionally, CD64/CD16A cDNA was inserted into the pKT2 sleeping beauty transposon vector and used along with SB100X transposase for iPSC transduction, as previously described (14). The nucleotide sequences of all constructs were confirmed by direct sequencing from both directions on an ABI 377 sequencer with ABI BigDye terminator cycle sequencing kit (Applied Biosystems, Foster City, CA).

### Cells

Fresh human peripheral blood leukocytes from plateletpheresis were purchased from Innovative Blood Resources (St. Paul, MN). Peripheral blood mononuclear cells were further enriched using Ficoll-Paque Plus (GE Healthcare Bio-Sciences AB, Uppsala, Sweden) and NK cells were purified by negative depletion using an EasySep human NK cell kit (StemCell Technologies, Cambridge, MA), as per the manufacturer’s instructions, with > 95% viability and 90–95% enrichment of CD56^+^CD3^−^ lymphocytes. Viable cell counting was performed using a Countess II automated cell counter (Life Technologies Corporation, Bothell, WA). The human NK cell line NK92 and the ovarian cancer cell line SKOV-3 were obtained from ATCC (Manassas, VA) and cultured per the manufacturer’s directions. The NK92 cells required IL-2 for growth (500 IU/ml), which was obtained from R&D Systems and the National Cancer Institute, Biological Resources Branch, Preclinical Biologics Repository (Frederick, MD). For all assays described below, cells were used when in log growth phase.

The iPSCs UCBiPS7, derived from umbilical cord blood CD34 cells, have been previously characterized and were cultured and differentiated into hematopoietic progenitor cells as described with some modifications (14, 28–31). iPSCs culture and hematopoietic differentiation was performed using TeSR-E8 medium and a STEMdiff Hematopoietic Kit (StemCell Technologies), which did not require the use of mouse embryonic fibroblast feeder cells, TrypLE adaption, spin embryoid body formation, or CD34^+^ cell enrichment. iPSC cells during passage were dissociated with Gentle Cell Dissociation Reagent (StemCell Technologies), and iPSC aggregates ≥ 50 µm in diameter were counted with a hemocytometer and diluted to 20 – 40 aggregates/ml with TeSR-E8 medium. Each well of a 12-well plate was pre-coated with Matrigel Matrix (Corning Inc., Tewksbury, MA) and seeded with 40 – 80 aggregates in 2 ml of TeSR-E8 medium. Cell aggregates were cultured for 24 hours before differentiation with the STEMdiff Hematopoietic Kit, as per the manufacturer’s instructions. At day 12 of hematopoietic progenitor cell differentiation, the percentage of hematopoietic progenitor cells was determined using flow cytometric analysis with anti-CD34, anti-CD45, and anti-CD43 mAbs. NK cell differentiation was performed as previously described (32). The iPSC-derived NK cells (referred to here as iNK cells) were expanded for examination using γ-irradiated K562-mbIL21–41BBL feeder cells (1:2 ratio) in cell expansion medium [60% DMEM, 30% Ham’s F12, 10% human AB serum (Valley Biomedical, Winchester, VA), 20µM 2-mercaptoethanol, 50 µM ethanolamine, 20 µg/ml ascorbic Acid, 5 ng/ml sodium selenite, 10 mM HEPES, and 100–250 IU/ml IL-2 (R&D Systems)], as described previously (14, 29–31)..

### Cell staining, flow cytometry and ELISA

For cell staining, nonspecific antibody binding sites were blocked and cells were stained with the indicated antibodies and examined by flow cytometry, as previously described (11, 14, 33). For controls, fluorescence minus one was used as well as appropriate isotype-matched antibodies since the cells of interest expressed FcRs. An FSC-A/SSC-A plot was used to set an electronic gate on leukocyte populations and an FSC-A/FSC-H plot was used to set an electronic gate on single cells. A Zombie viability kit was used to assess live vs. dead cells, as per the manufacturer’s instructions (BioLegend).

To assess the capture of soluble trastuzumab, rituximab, cetuximab, or L-selectin/Fc chimera, transduced NK cells were incubated with 5 μg/ml of antibody for 2 hours at 37°C in MEM-α basal media (Thermo Fisher Scientific, Waltham, MA) supplemented with IL-2 (200 IU/ml), HEPES (10mM), and 2-mercaptoethanol (0.1 mM), washed with MEM-α basal media, and then stained on ice for 30 minutes with a 1:200 dilution of APC-conjugated F(ab’)_2_ donkey anti-human IgG (H+L). To detect recombinant human L-selectin/Fc binding, cells were stained with the anti-L-selectin mAb APC-conjugated Lam1–116.

To compare CD16A and CD64/16A staining levels on NK92 cells, the respective transductants were stained with a saturating concentration of unconjugated anti-CD16 (3G8) or anti-CD64 (10.1) mAbs (5μg/ml), washed extensively with dPBS (USB Corporation, Cleveland, OH) containing 2% goat serum and 2mM sodium azide, and then stained with APC-conjugated F(ab’)_2_ goat anti-mouse IgG (H+L). This approach was used since directly conjugated anti-CD16 and anti-CD64 mAbs can vary in their levels of fluorophore labeling. ELISA was performed by a cytometric bead-based Flex Set assay to quantify human IFNγ levels (BD Biosciences, San Jose, CA), per the manufacturer’s instructions. All flow cytometric analyses were performed on a FACSCelesta instrument (BD Biosciences). Data was analyzed using FACSDIVA v8 (BD Biosciences) and FlowJo v10 (Ashland, OR).

### Cell-cell conjugation assay and ADCC

The pBMN-IRES-EGFP vector used to express CD64/16A in NK92 cells also expresses eGFP. These cells were incubated for 2 hours at 37°C in MEM-α basal media (Thermo Fisher Scientific, Waltham, MA) supplemented with IL-2 (200 IU/ml), HEPES (10mM), and 2-mercaptoethanol (0.1 mM). SKOV-3 cells were labeled with CellTrace Violet (Molecular Probes, Eugene, OR) per the manufacturer’s instructions, incubated with 5μg/ml trastuzumab for 30 minutes and washed with the MEM-α basal media. NK92 cells and SKOV-3 cells were then resuspended in the supplemented MEM-α basal media at 1×10^6^ and 2×10^6^/ml, respectively. For a 1:2 Effector:Target (E:T) ratio, 100μl of each cell type was mixed together, centrifuged for 1 minute at 20×g and incubated at 37°C for the indicated time points. After each time point, the cells were gently vortexed for 3 seconds and immediately fixed with 4°C 1% paraformaldehyde in dPBS. Samples were immediately analyzed by flow cytometry. The percentage of conjugated NK cells was calculated by gating on eGFP and CellTrace Violet double-positive events.

To evaluate ADCC, a DELFIA EuTDA-based cytotoxicity assay was used according to the manufacturer’s instructions (PerkinElmer, Waltham, MA). Briefly, target cells were labeled with Bis(acetoxymethyl)-2–2:6,2 terpyridine 6,6 dicarboxylate (BATDA) for 30 minutes in their culture medium, washed in culture medium, and pipetted into a 96-well non-tissue culture-treated U-bottom plates at a density of 8×10^4^ cells/well. A tumor targeting mAb was added at the indicated concentrations of 5μg/mL and NK cells were added at the indicated E:T ratios. The plates were centrifuged at 400×g for 1 minute and then incubated for 2 hours in a humidified 5% CO_2_ atmosphere at 37°C. At the end of the incubation, the plates were centrifuged at 500×g for 5 minutes and supernatants were transferred to a 96 well DELFIA Yellow Plate (PerkinElmer) and combined with europium. Fluorescence was measured by time-resolved fluorometry using a BMG Labtech CLARIOstar plate reader (Cary, NC). BATDA-labeled target cells alone with or without therapeutic antibodies were cultured in parallel to assess spontaneous lysis and in the presence of 2% Triton-X to measure maximum lysis. The level of ADCC for each sample was calculated using following formula: 
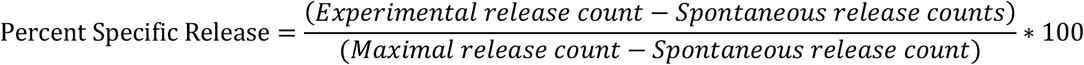
For each experiment, measurements were conducted in triplicate using three replicate wells.

### Statistical analyses

Statistical analyses were performed by use of GraphPad Prism (GraphPad Software, La Jolla, CA, USA). After assessing for approximate normal distribution, all variables were summarized as mean ± SD. Comparison between 2 groups was done with Student’s t-test, with p<0.05 taken as statistically significant.

## Results

### Expression and function of CD64/16A in NK92 cells

We engineered a recombinant FcγR that consists of the extracellular region of human CD64 and the transmembrane and cytoplasmic regions of human CD16A, referred to as CD64/16A (**Fig. 1A**). The recombinant receptor was stably expressed in the human NK cell line NK92. These cells lack endogenous FcγRs but transduced cells expressing exogenous CD16A can mediate ADCC (14, 20, 27). As shown is **Figure 1B**, an anti-CD64 mAb stained NK92 cells expressing CD64/16A cells, but not parent NK92 cells or NK92 cells expressing CD16A. An anti-CD16 mAb stained NK92 cells expressing CD16A, but not NK92 cells expressing CD64/16A or parent NK92 cells (**Fig. 1B**). CD16A undergoes ectodomain shedding by ADAM17 upon NK cell activation resulting in its rapid downregulation in expression (10–13, 33). CD16A and its isoform CD16B on neutrophils is cleaved by ADAM17 (10), and this occurs at an extracellular region proximal to the cell membrane (13, 14). The ADAM17 cleavage region of CD16A is not present in CD64 or CD64/16A (**Fig. 1A**). We found that CD16A underwent a > 50% decrease in expression upon NK92 stimulation by ADCC, whereas CD64/16A demonstrated little to no downregulation (**Fig. 1C**).

**Figure 1.**
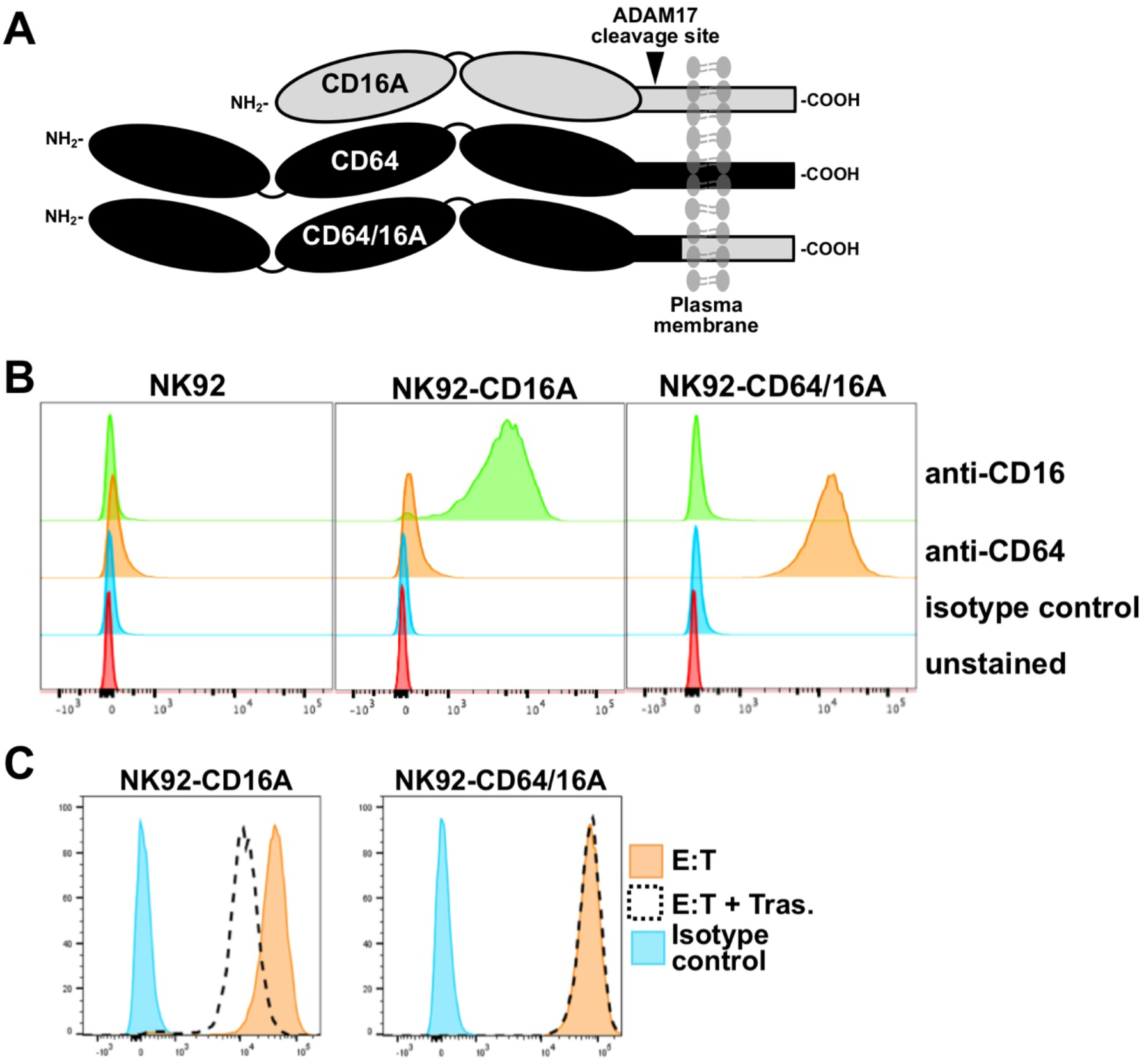
Expression of CD64/16A by NK92 cells. **(A)** Schematic representation of the cell membrane forms of CD16A, CD64, and CD64/16A. CD16A undergoes ectodomain shedding by ADAM17 at a membrane proximal location, as indicated, which is not present in CD64 and CD64/16A. **(B)** NK92 parental cells, NK92-CD16A cells, and NK92-CD64/16A cells were stained with an anti-CD16, anti-CD64, or an isotype-matched negative control mAb and examined by flow cytometry. **(C)** NK92-CD16A and NK92-CD64/16A cells were incubated with SKOV-3 cells with or without trastuzumab (5μg/ml) at 37°C (E:T = 1:1) for 2 hours. The NK92-CD16A and NK92-CD64/16A cells were then stained with an anti-CD16 mAb or an anti-CD64 mAb, respectively, and examined by flow cytometry. Nonspecific antibody labeling was determined using the appropriate isotype-negative control mAb. Data is representative of at least 3 independent experiments.

To evaluate the function of CD64/16A, we examined its ability to initiate E:T conjugation, induce ADCC, and stimulate IFN-g production upon NK cell engagement of antibody-bound tumor cells. Prior to the release of its granule contents, an NK cell must form a stable conjugate with a target cell. Using a two-color flow cytometric approach, we examined the conjugation of NK92-CD64/16A cells and SKOV-3 cells, an ovarian cancer cell line that expresses HER2. This assay was performed in the absence and presence of the anti-HER2 therapeutic mAb trastuzumab. The bicistronic vector containing CD64/16A also expressed eGFP and its fluorescence was used to identify the NK92 cells. SKOV-3 cells were labeled with the fluorescent dye CellTrace Violet. E:T conjugation resulted in two-color events that were enumerated by flow cytometry. The incubation of NK92-CD64/16A cells with SKOV-3 cells resulted in a very low level of conjugation after initial exposure that increased after 60 minutes of exposure (**Fig. 2A**). However, in the presence of trastuzumab, NK92-CD64/16A cell and SKOV-3 conjugation was appreciably enhanced (**Fig. 2A**). This increase in conjugation corresponded with higher levels of target cell killing. As shown in **Figure 2B**, SKOV-3 cell cytotoxicity by NK92-CD64/16A cells varied depending on the trastuzumab concentration and E:T ratio. To confirm the role of CD64/16A in the induction of target cell killing, we performed the ADCC assay in the presence and absence of the anti-CD64 mAb 10.1 (**Fig. 2C**), which blocks IgG binding (34). Cytokine production is also induced during ADCC and NK cells are major producers of IFNγ (4, 35). NK92-CD64/16A cells exposed to SKOV-3 cells and trastuzumab produced considerably higher levels of IFNγ than when exposed to SKOV-3 cells alone (**Fig. 2D**). Taken together, the above findings demonstrate that the CD64 component of the recombinant receptor engages tumor-bound antibody, and that the CD16A component promotes intracellular signaling leading to degranulation and cytokine production.

**Fig. 2.**
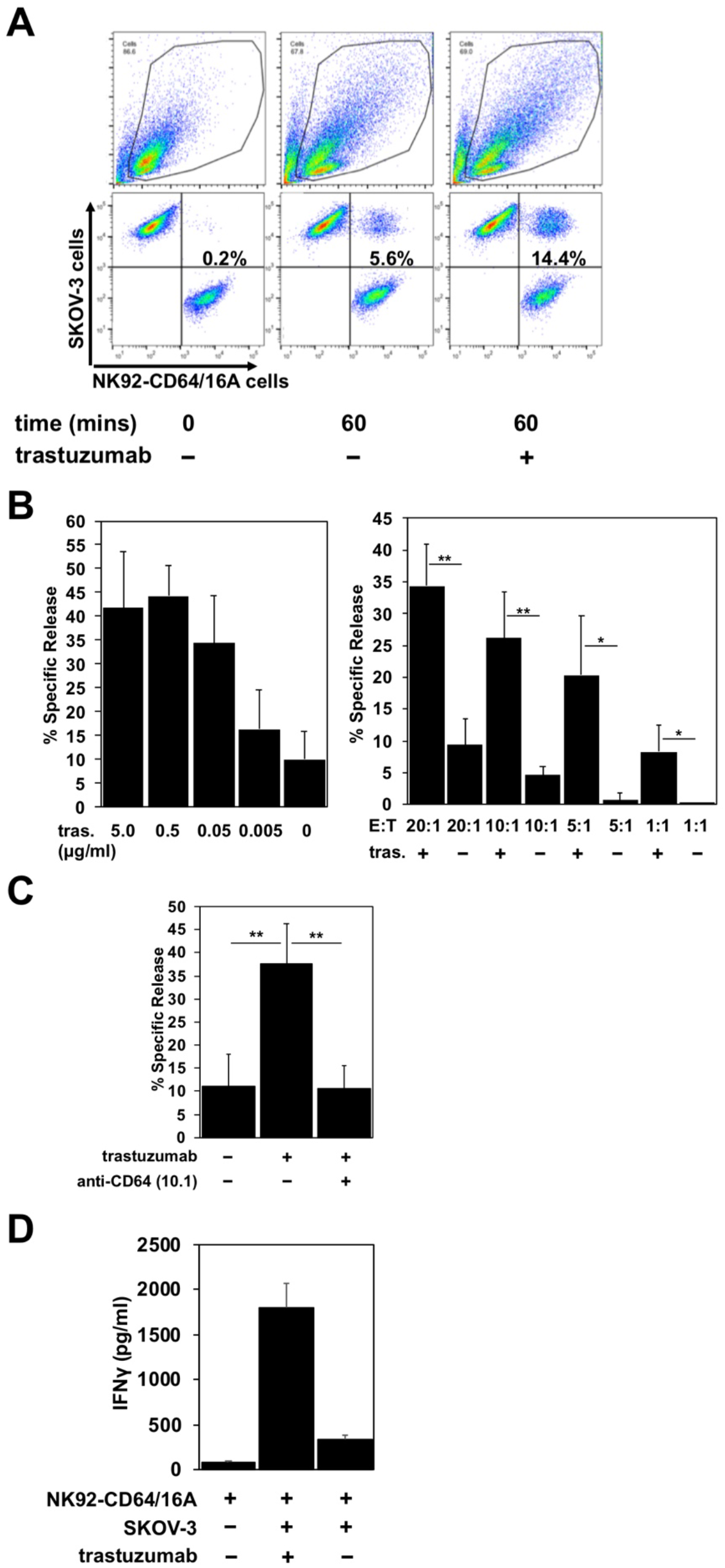
CD64/16A promotes target cell conjugation, ADCC, and IFNγ production. **(A)** NK92-CD64/16A cells expressing eGFP and SKOV-3 cells labeled CellTrace Violet were mixed at an E:T ratio of 1:2 with or without trastuzumab (5μg/ml), incubated at 37 °C for 60 minutes, fixed, and then analyzed by flow cytometry, as described in the Materials and Methods. Representative data from at least three independent experiments are shown. **(B)** NK92-CD64/16A cells were incubated with SKOV-3 cells (E:T = 20:1) and trastuzumab (tras.) at the indicated concentrations (left panel), or with SKOV-3 cells at the indicated E:T ratios in the presence or absence of trastuzumab (5μg/ml) (right panel) for 2 hours at 37 °C. Data are represented as % specific release and the mean±SD of 3 independent experiments is shown. Statistical significance is indicated as *p<0.05, **p<0.01. **(C)** NK92-CD64/16A cells were incubated with SKOV-3 cells (E:T = 20:1) in the presence or absence of trastuzumab (5μg/ml) and the anti-CD64 mAb 10.1 (10μg/ml), as indicated, for 2 hours at 37 °C. Data are represented as % specific release and the mean±SD of 3 independent experiments is shown. Statistical significance is indicated as **p<0.01. **(D)** NK92-CD64/16A cells were incubated with SKOV-3 cells (E:T = 1:1) with or without trastuzumab (5μg/ml) for 2 hours at 37 °C. Secreted IFNγ levels were quantified by ELISA. Data is shown as mean of 2 independent experiments.

CD64 is distinguished from the other FcγR members by its unique third extracellular domain, which contributes to its high affinity and stable binding to soluble monomeric IgG (26). We compared the ability of NK92 cells expressing CD64/16A or the CD16A-176V higher affinity variant to capture soluble therapeutic mAbs. The NK92 cell transductants examined expressed similar levels of CD64/16A and CD16A (**Figure 3A**). NK92 cell transductants were incubated with trastuzumab for 2 hours, excess antibody was washed away, stained with a fluorophore-conjugated anti-human IgG antibody, and then evaluated by flow cytometry. As shown in **Figure 3B**, NK92-CD64/16A cells captured considerably higher levels of trastuzumab than did the NK92-CD16A cells (8.1 fold increase ± 1.3, mean±SD of 3 independent experiments). Moreover, the NK92-CD64/16A cells efficiently captured the tumor-targeting mAbs Erbitux/cetuximab and Rituxan/rituximab, as well as the fusion protein L-selectin/Fc (**Fig. 3C**). We then tested whether NK92-CD64/16A cells with a captured tumor-targeting mAb mediated ADCC. For this assay, equal numbers of NK92-CD64/16A and NK92-CD16A cells were incubated with the same concentration of soluble trastuzumab, washed, and exposed to SKOV-3 cells. Target cell killing by NK92-CD64/16A cells with captured trastuzumab was significantly higher than NK92-CD64/16A cells alone, and was far superior to NK92-CD16A cells treated with or without trastuzumab at all E:T ratios examined (**Fig. 3D**). In contrast, SKOV-3 cytotoxicity by NK92-CD16A and NK92-CD64/16A cells was not significantly different if trastuzumab was present and not washed out (**Fig. 3E**), thus demonstrating equivalent cytotoxicity by both transductants. Taken together, these findings show that NK92 cells expressing CD64/16A can stably bind soluble anti-tumor mAbs and IgG fusion proteins, and that these can serve as targeting elements to kill cancer cells.

**Fig. 3.**
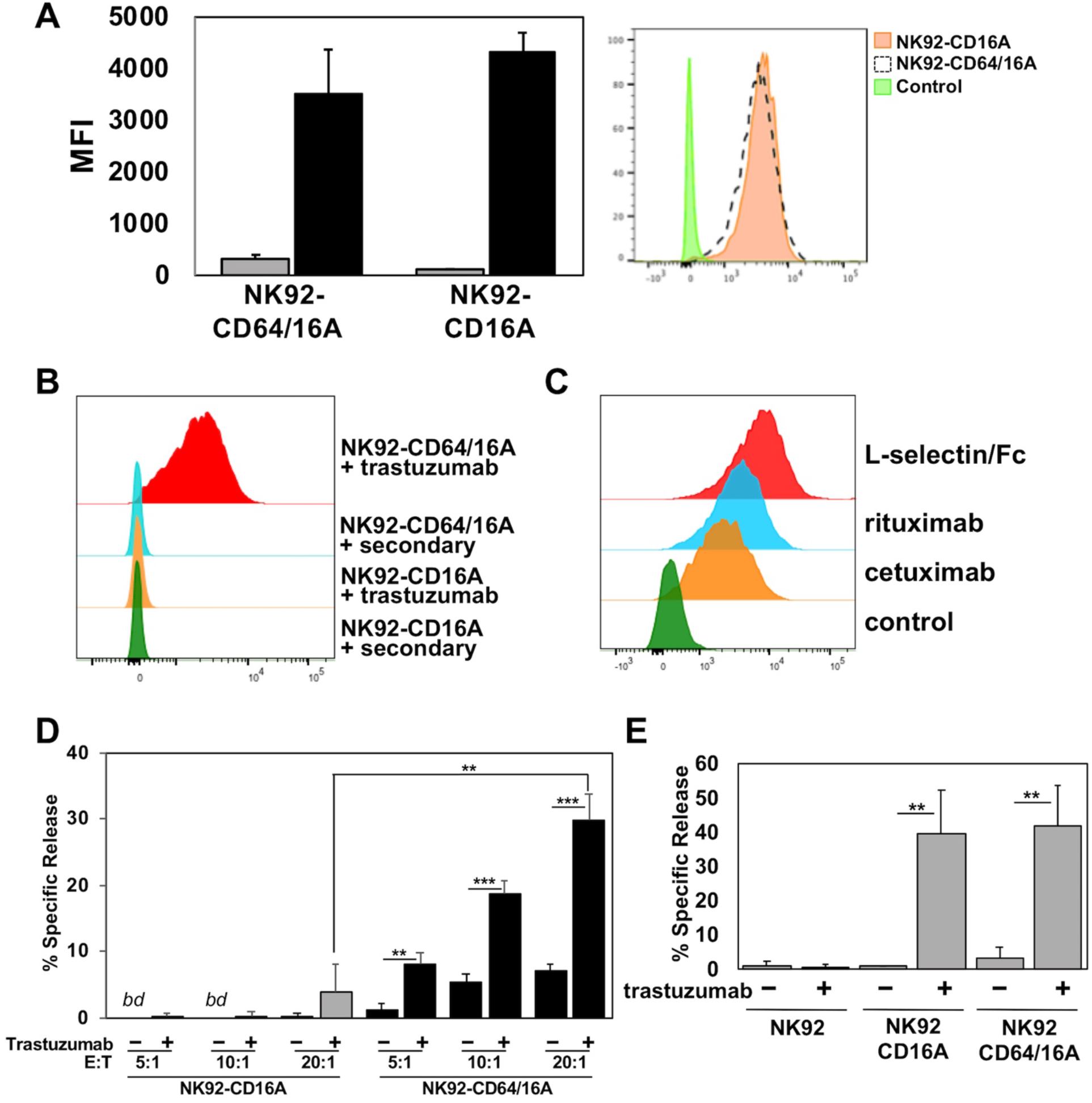
CD64/16A attaches to soluble tumor-targeting mAbs and IgG fusion proteins. **(A)** Relative expression levels of CD16A and CD64/16A on NK92 cells were determined by cell staining with anti-CD16 and anti-CD64 mAbs (black bars), respectively, or an isotype-matched negative control antibody (gray bars). The bar graph shows mean fluorescence intensity (MFI)±SD of three independent experiments. Representative flow cytometric data are shown in the histogram overlay. The dashed line histogram shows CD64 staining of NK92-CD64/16A cells, the orange-filled histogram shows CD16A staining of NK92-CD16A cells, and the green-filled histogram shows isotype control antibody staining of the NK92-CD16A cells. **(B)** NK92-CD16A and NK92-CD64/16A cells were incubated with or without trastuzumab (5μg/ml) for 2 hours at 37°C, washed, stained with a fluorophore-conjugated anti-human secondary antibody, and analyzed by flow cytometry. Data is representative of at least 3 independent experiments. **(C)** NK92-CD64/16A cells were incubated with cetuximab or rituximab (5μg/ml for each), washed, and then stained with a fluorophore-conjugated anti-human secondary antibody. Control represents cells stained with the anti-human secondary antibody only. NK92-CD64/16A cells were also incubated with L-selectin/Fc (5μg/ml), washed, and then stained with a fluorophore-conjugated anti-L-selectin mAb. NK92 cells lack expression of endogenous L-selectin (data not shown). All staining was analyzed by flow cytometry. Data shown are representative of 3 independent experiments. **(D)** NK92-CD16A and NK92-CD64/16A cells were incubated in the presence or absence of trastuzumab (5μg/ml), washed, and exposed to SKOV-3 cells at the indicated E:T cell ratios for 2 hours at 37°C. Data is shown as mean±SD of 3 independent experiments. Statistical significance is indicated as **p<0.01, ***p<0.001. *bd* = below detection, (i.e., < spontaneous release by negative control cells). **(E)** NK92-CD16A and NK92-CD64/16A cells were incubated with SKOV-3 cells (E:T = 10:1) in the presence or absence of trastuzumab (5μg/ml), as indicated, for 2 hours at 37°C. Data is shown as mean±SD of 3 independent experiments. Statistical significance is indicated as **p<0.01.

### Expression and function of CD64/16A in iPSC-derived NK cells

We also examined the function of CD64/16A in engineered primary NK cells. Genetically modifying peripheral blood NK cells by retroviral or lentiviral transduction at this point has been challenging (36). Embryonic stem cells and iPSCs can be differentiated into cytolytic NK cells *in vitro* (28–31, 37), and these cells are highly amendable to genetic engineering (14, 30, 38, 39). Undifferentiated iPSCs were transduced to express CD64/16A using a sleeping beauty transposon plasmid for nonrandom gene insertion and stable expression. iPSCs were differentiated into hematopoietic cells and then iNK cells by a two-step process (28, 29). For this study, we modified the hematopoietic differentiation method to streamline the procedure by using a commercially available media and hematopoietic differentiation kit, as described in the Materials and Methods. CD34^+^CD43^+^CD45^+^ cells were generated, further differentiated into iNK cells, and these cells were expanded for analysis using recombinant IL-2 and aAPCs. CD56^+^CD3^−^ is a hallmark phenotype of human NK cells, and these cells composed the majority of our differentiated cell population (**Fig. 4**). We also assessed the expression of a number of activating and inhibitory receptors on the iNK cells and compared this to peripheral blood NK cells. Certain receptors were expressed by similar proportions of the two NK cell populations, such as CD16A; however, the expanded iNK cells did lack expression of the inhibitory KIR receptors KIR2DL2/3, KIR2DL1, and KIR3DL1 and also certain activating receptors (NKp46 and NKG2D) (**Fig. 4**). Another difference compared to peripheral blood NK cells was that the iNK cells were stained with an anti-CD64 mAb (**Fig. 4**), demonstrating the expression of CD64/16A.

**Fig. 4.**
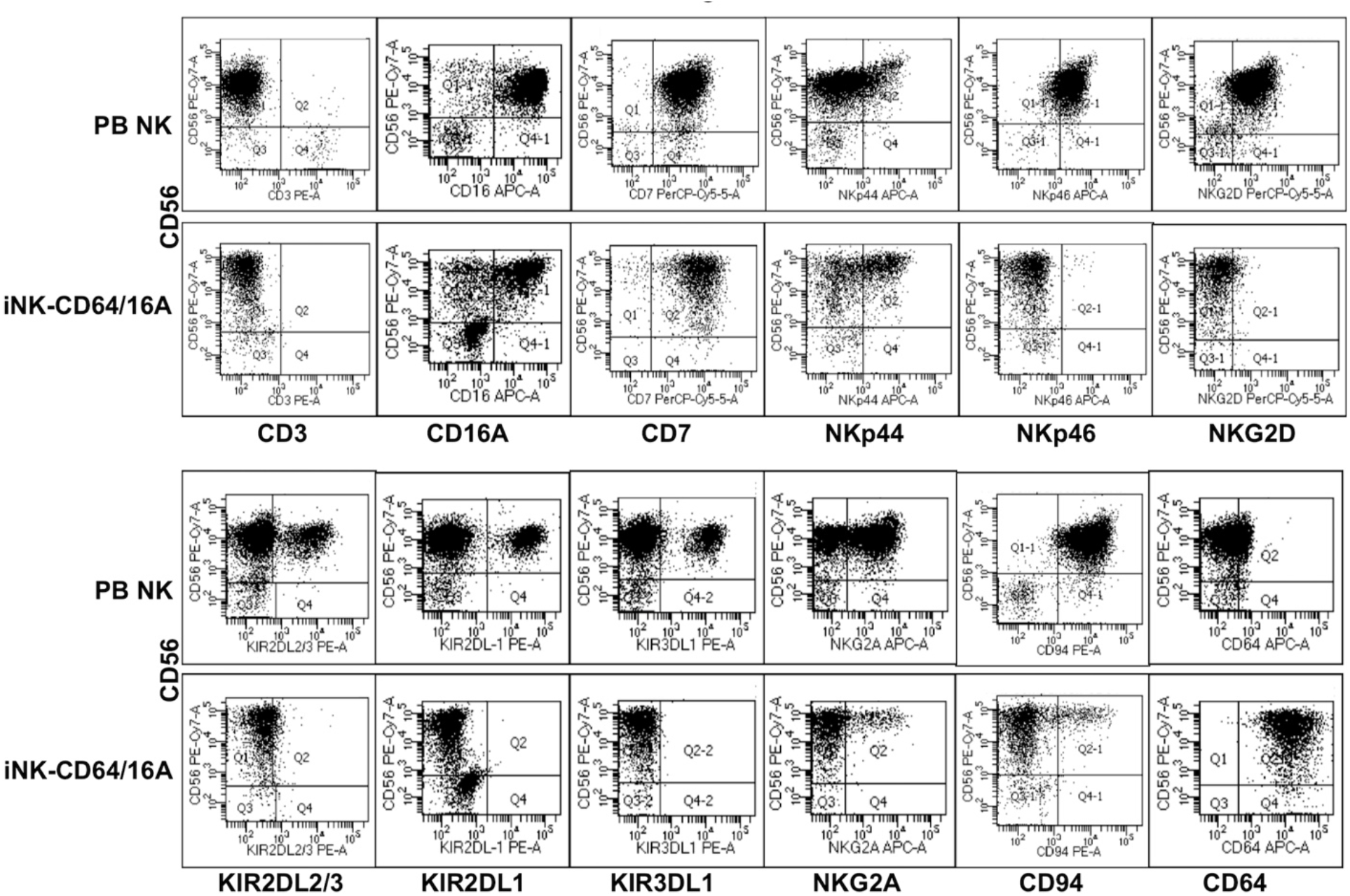
Generation of iNK cells expressing CD64/CD16A. iPSCs were transduced to stably express CD64/16A, differentiated into NK cells, and then expanded using K562-mbIL21–41BBL feeder cells, as described in the Materials and Methods. iNK-CD64/16A cells and freshly isolated peripheral blood (PB) NK cells enriched from adult peripheral blood were stained for CD56, CD3 and various inhibitory and activating receptors, as indicated. CD64/16A expression was determined by staining the cells with an anti-CD64 mAb. Representative data from at least three independent experiments are shown.

To assess the function of CD64/16A in iNK cells, we compared iNK cells derived from iPSCs transduced with either an pKT2 empty vector or pKT2-CD64/16A. The NK cell markers mentioned above were expressed at similar levels and proportions by two iNK cell populations (data not shown), including CD16A (**Fig. 5A**), but only iNK-CD64/16A cells were stained by an anti-CD64 mAb (**Fig. 5A**). Both iNK tranductants demonstrated increased SKOV-3 cell killing when in the presence of trastuzumab, yet iNK-CD64/16A cells mediated significantly higher levels of ADCC than did the iNK-pKT2 control cells (**Fig. 5B**). The anti-CD16 function blocking mAb 3G8, but not the anti-CD64 mAb 10.1, effectively inhibited ADCC by the iNK-pKT2 cells (**Fig. 5B**). Conversely, 10.1, but not 3G8, blocked ADCC by the iNK-CD64/16A cells (**Fig. 5B**). These findings show that the iNK cells were cytolytic effectors responsive to CD16A and CD64/16A engagement of antibody-bound tumor cells. We also treated iNK-CD64/16A and iNK-pKT2 cells with soluble trastuzumab, washed away excess antibody, and exposed them to SKOV-3 cells. Under these conditions, ADCC by the iNK-CD64/16A cells was strikingly higher than the iNK-pKT2 cells (**Fig. 5C**), further establishing that CD64/16A can capture soluble anti-tumor mAbs that serve as a targeting element for tumor cell killing.

**Fig 5.**
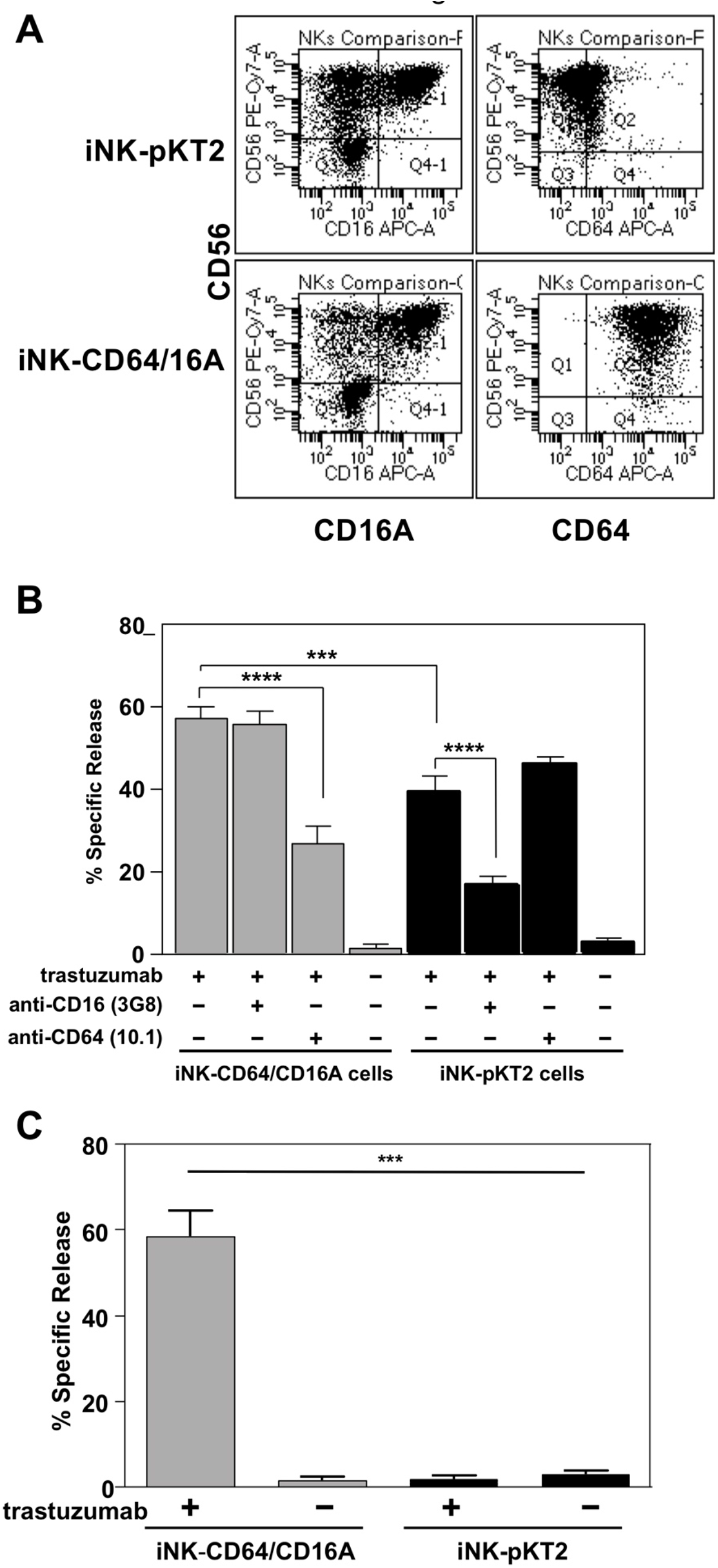
iNK-CD64/16A cells show enhanced ADCC compared to iNK-pKT2 control cells. (**A)** NK cells derived from empty vector (iNK-pKT2) or CD64/16A (iNK-CD64/16A) transduced iPSCs were stained for CD56, CD64, and CD16A, as indicated. **(B)** iNK-pKT2 and iNK-CD64/16A cells were incubated with SKOV-3 cells (E:T = 10:1) in the presence or absence of trastuzumab (5μg/ml), the function blocking anti-CD16 mAb 3G8 (5μg/ml), and the function blocking anti-CD64 mAb 10.1 (5μg/ml), as indicated, for 2 hours at 37°C. Data is shown as mean±SD of 3 independent experiments. Statistical significance is indicated as ***p<0.001; ****p<0.0001. **(C)** iNK-pKT2 and iNK-CD64/16A cells were incubated in the presence or absence of trastuzumab (5μg/ml), washed, and exposed to SKOV-3 cells (E:T = 10:1) for 2 hours at 37°C. Data is shown as mean±SD of 3 independent experiments. Statistical significance is indicated as ***p<0.001.

## Discussion

CD16A has an exclusive role in inducing ADCC by human NK cells (2–4). The affinity of antibody binding and the expression levels of this IgG Fc receptor modulate NK cell effector functions and affect the efficacy of tumor-targeting therapeutic mAbs (4, 11, 19, 20). To enhance anti-tumor antibody binding by NK cells, we expressed a novel recombinant FcγR consisting of the extracellular region of the high affinity FcγR CD64 and the transmembrane and intracellular regions of CD16A. NK cells expressing CD64/16A facilitated cell conjugation with antibody-bound tumor cells, cytotoxicity, and IFNγ production, demonstrating function by both components of the recombinant FcγR. CD64/16A lacks the ADAM17 cleavage region found in CD16A and it did not undergo the same level of downregulation in expression during ADCC. Consistent with the ability of CD64 to stably bind soluble monomeric IgG, NK cells expressing CD64/16A could capture soluble anti-tumor therapeutic mAbs and kill target cells.

CD64/16A was shown to be functional in two NK cell platforms, the NK92 cell line and primary NK cells derived from iPSCs. NK92 cells lack inhibitory KIR receptors and show high levels of natural cytotoxicity compared to other NK cell lines derived from patients (40). NK92 cells have been broadly used to express modified genes to direct their cytolytic effector function, have been evaluated in preclinical studies, and are undergoing clinical testing in cancer patients (40, 41). iPSCs are also very amendable to genetic engineering and can be differentiated into NK cells expressing various receptors to direct their effector functions (14, 30, 38, 39). The iNK cells generated in this study lacked several inhibitory and activating receptors indicating an immature state. In previous studies we have generated iNK cells with a phenotype indicative of a more mature cellular stage (29–31), which may be the result of different culture conditions. A key change for the current study was the use of a hematopoietic differentiation kit to simplify the differentiation procedure. The phenotype of the iNK cells will be important for the desired effector functions. It could be beneficial if therapeutic iNK cells administered to cancer patients lacked inhibitory receptors and certain activating receptors in order to direct and optimize their tumor cell killing by engineered receptors. The iNK cells did express endogenous CD16A and mediated ADCC, thus they were cytotoxic effector cells. We found that for pKT2 vector control iNK cells, ADCC was blocked by an anti-CD16 mAb. Interestingly, ADCC by the iNK-CD64/16A cells was blocked by an anti-CD64 mAb but not by an anti-CD16 mAb. Why endogenous CD16A in the iNK-CD64/16A cells did not have a role in the *in vitro* ADCC assay is unclear at this time. This may be due to a competitive advantage by CD64/16A in binding antibody and/or in utilizing the same pool of downstream signaling factors.

An individual NK cell can kill multiple tumor cells in different manners. This includes by a process of sequential contacts and degranulations, referred to as serial killing (42, 43), and by the localized dispersion of its granule contents that kills surrounding tumor cells, referred to as bystander killing (44). Further studies are required to determine the effects of CD64/16A expression on these killing processes during ADCC. Inhibiting CD16A shedding has been reported to slow NK cell detachment from target cells and reduce serial killing by NK cells *in vitro* (45). Due to the CD64 component and its lack of ectodomain shedding, NK cells expressing CD64/16A could be less efficient at serial killing and more efficient at bystander killing. An important next step will be to assess the anti-tumor activity of NK cells expressing CD64/16A *in vivo*, and the studies will include the use of NK92-CD64/16A cells and iNK-CD64/16A cells in tumor xenograft models.

Therapeutic mAbs have become one of the fastest growing classes of drugs, and tumor-targeting mAbs are the most widely used and characterized immunotherapy for hematologic and solid tumors (46). NK cells expressing CD64/16A have several potential advantages as a combination therapy, as their capture of anti-tumor mAbs, either individually or in combination, prior to adoptive transfer provides diverse options for switchable targeting elements. Modifying NK cells expressing CD64/16A with an antibody would also reduce the dosage of therapeutic antibodies administered to patients. We showed that fusion proteins containing a human IgG Fc region, such as L-selectin/Fc, can also be captured by CD64/16A and this may provide further options for directing the tissue and tumor antigen targeting of engineered NK cells. Advantages of the NK92 and iNK cell platforms for adoptive cell therapies is that they can be readily gene modified on a clonal level and expanded into clinical-scalable cell numbers to produce engineered NK cells with improved effector activities as an off-the-shelf therapeutic for cancer immunotherapy (37, 38, 40, 41, 47).

## Author Contributions

BW and JW collected, assembled, analyzed and interpreted the data, and wrote the manuscript. KS collected, analyzed, and interpreted the data, and revised the manuscript. RH, HM, DM, YL, and AR collected, analyzed, and interpreted the data. DK analyzed the data and revised the manuscript. All authors contributed to manuscript preparation, read, and approved the submitted version.

